# The Fruit Fly Brain Observatory: from structure to function

**DOI:** 10.1101/092288

**Authors:** Nikul H. Ukani, Chung-Heng Yeh, Adam Tomkins, Yiyin Zhou, Dorian Florescu, Carlos Luna Ortiz, Yu-Chi Huang, Cheng-Te Wang, Paul Richmond, Chung-Chuan Lo, Daniel Coca, Ann-Shyn Chiang, Aurel A. Lazar

**Affiliations:** Department of Electrical Engineering, Columbia University, New York, NY 10027, USA; Department of Automatic Control & Systems Engineering, The University of Sheffield, Sheffield, S1 3JD, UK; Brain Research Center, National Tsing Hua University, Hsinchu 30013, Taiwan; Institute of Systems Neuroscience, National Tsing Hua University, Hsinchu 30013, Taiwan; Department of Life Science, National Tsing Hua University, Hsinchu 30013, Taiwan; Department of Computer Science, The University of Sheffield, Sheffield, S1 4DP, UK; Genomics Research Center, Academia Sinica, Nankang, Taipei 11529, Taiwan; Institute of Physics, Academia Sinica, Nankang, Taipei 11529, Taiwan; Department of Biomedical Science and Environmental Biology, Kaohsiung Medical University, Kaohsiung 80708, Taiwan; Kavli Institute for Brain and Mind, University of California, San Diego, La Jolla, California 92093, USA

## Abstract

The Fruit Fly Brain Observatory (FFBO) is a collaborative effort between experimentalists, theorists and computational neuroscientists at Columbia University, National Tsing Hua University and Sheffield University with the goal to (i) create an open platform for the emulation and biological validation of fruit fly brain models in health and disease, (ii) standardize tools and methods for graphical rendering, representation and manipulation of brain circuits, (iii) standardize tools for representation of fruit fly brain data and its abstractions and support for natural language queries, (iv) create a focus for the neuroscience community with interests in the fruit fly brain and encourage the sharing of fruit fly brain structural data and executable code worldwide. NeuroNLP and NeuroGFX, two key FFBO applications, aim to address two major challenges, respectively: i) seamlessly integrate structural and genetic data from multiple sources that can be intuitively queried, effectively visualized and extensively manipulated, ii) devise executable brain circuit models anchored in structural data for understanding and developing novel hypotheses about brain function. NeuroNLP enables researchers to use plain English (or other languages) to probe biological data that are integrated into a novel database system, called NeuroArch, that we developed for integrating biological and abstract data models of the fruit fly brain. With powerful 3D graphical visualization, NeuroNLP presents a highly accessible portal for the fruit fly brain data. NeuroGFX provides users highly intuitive tools to execute neural circuit models with Neurokernel, an open-source platform for emulating the fruit fly brain, with full data support from the NeuroArch database and visualization support from an interactive graphical interface. Brain circuits can be configured with high flexibility and investigated on multiple levels, e.g., whole brain, neuropil, and local circuit levels. The FFBO is publicly available and accessible at http://fruitflybrain.org from any modern web browsers, including those running on smartphones.

## Additional Details

Given the enormous complexity of the brain, finding effective ways of collaboration is of great interest to the neuroscience research community [1]. We describe here a path to move forward by harnessing the energy and know-how of three research groups located on three continents.

## Architecture of the FFBO

Supporting the two FFBO web applications is a highly sophisticated software architecture sketched in Figure 1. The two key components at the back-end are NeuroArch [2] and Neurokernel [3]. NeuroArch is a novel, graph-based database that supports integration of biological and modeling data within a single unified database accessed via a query-based API designed with the explicit aim of facilitating the generation of executable neural circuit models. The use of a data representation and interface specifically designed to support construction of executable models distinguishes NeuroArch from other fruit fly biological databases and their associated query interfaces. This support for executable circuits enables seamless integration with Neurokernel, a platform for the emulation of the fruit fly brain, to execute circuits retrieved from NeuroArch, or alternative models that are algorithmically generated, e.g., using virtual genetic manipulation. In addition, the architecture includes a natural language processing (NLP) module that handles the parsing of plain English queries.

**Figure 1:**
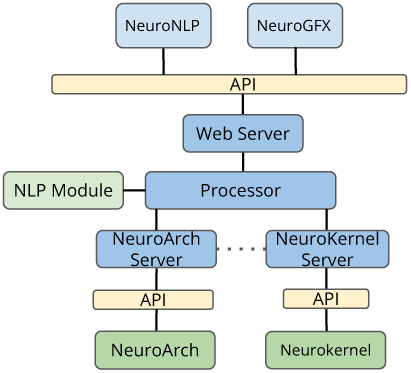
FFBO Architecture.

The FFBO thereby embraces a simple yet powerful workflow for investigating the fruit fly brain function. This workflow is shown in Figure 2. Biological data from multiple sources are integrated/aggregated into the NeuroArch database. Executable circuit models can be constructed from the structural data and executed by Neurokernel.

**Figure 2:**
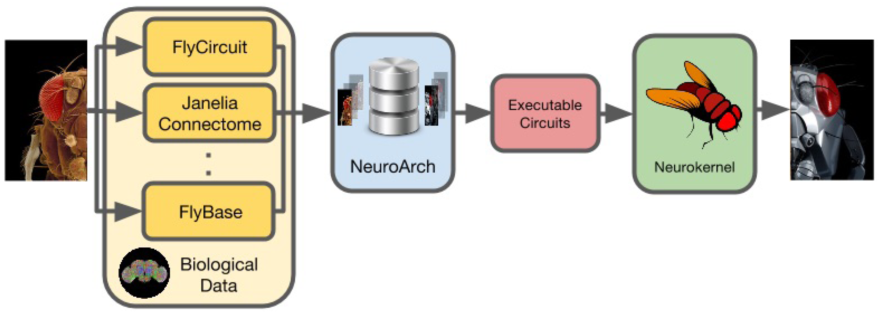
Workflow of the Fruit Fly Brain Observatory.

Currently NeuroArch stores fruit fly connectome data from two sources: the FlyCircuit database [4] and the Janelia Fly Medulla data [5]. The former hosts meso-scale connectome data on the whole-brain level, and the latter contains detailed, micro-scale data about exact chemical synaptic information between neurons in a limited region of the Medulla neuropil.

## NeuroNLP

In NeuroNLP, neurons in the NeuroArch databases can be queried based on the neuropil that they reside in, their innervation/connection pattern and their neurotransmitter type. For example, to query and visualize the glutamatergic local neurons in the antennal lobe, one can simply type the query: “show glutamatergic local neurons in AL”. A flexible GUI provides additional power to refine the control of circuit visualization. Extensive demos are provided for first time users. The user interface is shown in Figure 3 with an example showing 16 Lobula Plate Tangential Cells (LPTC).

**Figure 3:**
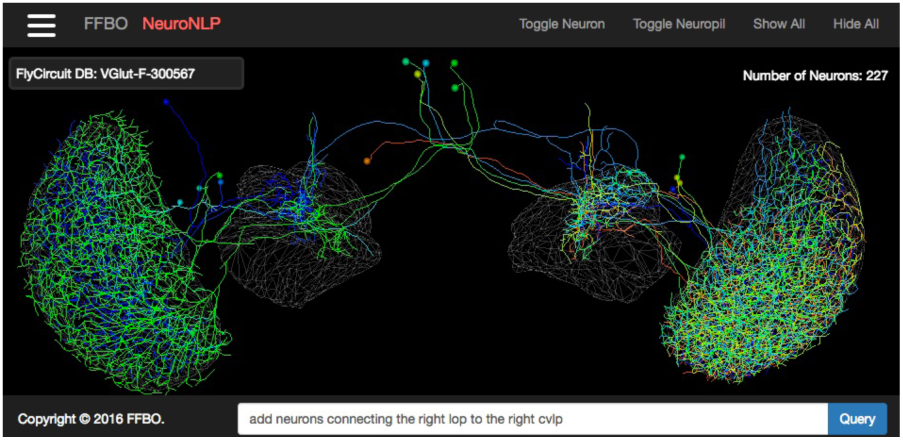
16 LPTCs extracted from the database using NeuroNLP.

## NeuroGFX

To bridge the gap between structure and function, structural information about the fruit fly brain should be mapped into executable brain circuit models. NeuroGFX provides a programming interface to i) query and retrieve modeling data from the NeuroArch database, ii) configure circuit diagrams side-by-side with morphological information, e.g., silencing certain neurons (see shaded orange block in Figure 4), iii) execute circuit on a Neurokernel server that can reside on a GPU server in a lab or on the Amazon Computing Cloud, and iv) visualize and playback results with complete flexibility of what to show.

**Figure 4:**
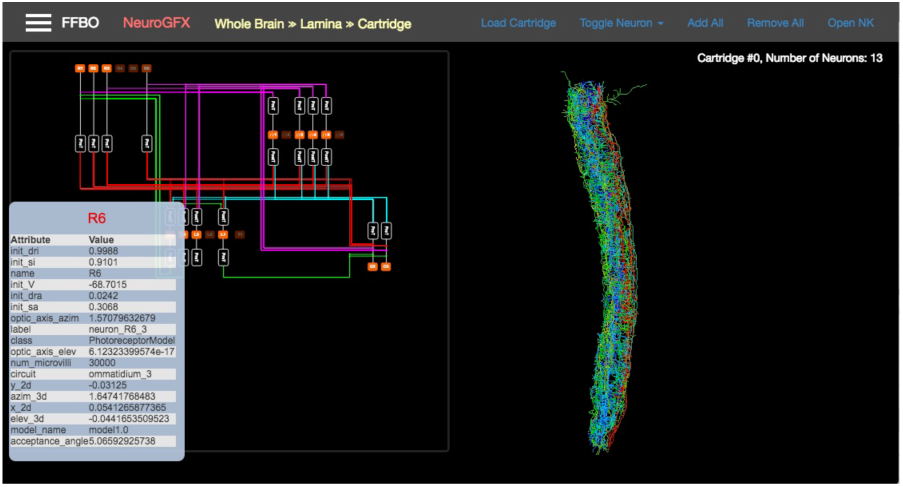
Exploring the structure and function of a lamina cartridge with NeuroGFX.

The FFBO can dramatically increase the pace of discovery by enabling neurobiologists and computational neuroscientists alike to generate and test new hypotheses, by interrogating, visualizing and simulating the fruit fly brain at different scales, from neural circuits to neuropils and ultimately the entire brain.

